# Mechanistic studies of autophagic cargo recruitment and membrane expansion through in vitro reconstitution

**DOI:** 10.1101/2024.12.24.630225

**Authors:** Wenxin Zhang, Thomas Litschel, Rocco D’Antuono, Colin Davis, Anne Schreiber, Sharon A. Tooze

## Abstract

Autophagy is a highly conserved catabolic pathway to remove deleterious cytosolic material to maintain cellular homeostasis and cell survival. Upon autophagy induction, a unique double-membraned structure, called a phagophore, forms and expands into a cup shape to engulf these cytosolic substrates. ATG8 proteins are covalently conjugated to autophagic membranes by lipidation of phosphatidylethanolamine (PE) and are thought to localise on both sides of the phagophore membrane. ATG8 conjugated on the inner membrane of the phagophore recruits autophagy cargo receptors, such as p62. To recapitulate events on the inner membrane, we used giant unilamellar vesicles (GUVs) as a model membrane and encapsulated proteins of interest inside GUVs, thus generating a membrane platform to which ATG8 proteins could be localised on the inner leaflet of the vesicles. We reconstituted WIPI2b-directed and cargo-directed ATG8 lipidation inside the GUVs and revealed distinct roles of WIPI2b and p62 in initiating the ATG conjugation cascade. Furthermore, we showed that p62 or p62 droplets were recruited to the inner membrane of the GUVs though interaction with membrane-bound ATG8s. Using a bead-based membrane expansion assay, we demonstrated a redistribution and local enrichment of membrane-bound ATG8s across the membrane upon interaction with p62 and p62 droplets. Our study provides novel model systems to investigate the interactions on the inner membrane of the phagophore and reveals fundamental molecular insights into phagophore membrane bending. This process is directed by ATG8-cargo interaction, during which cargo receptors concentrate ATG8 proteins on the inner surface of the phagophore membrane.

## Introduction

Macroautophagy (hereafter autophagy) is the main degradative pathway to deliver long-lived, redundant or damaged cytoplasmic materials to the lysosome (1, 2). Upon autophagy induction, a unique double membrane structure, the phagophore, is formed, growing into a cup-shaped structure which captures cytoplasmic components (3–5). A critical question pertinent to phagophore formation is how the membrane acquires its shape as it expands. The machinery, in particular the proteins and lipids, that direct the shaping are not fully understood.

It has been proposed that phagophore membrane bending is directed by cargo-ATG8 interactions (6, 7). ATG8 is a ubiquitin-like protein covalently conjugated to phospholipid (PE or PS) on phagophore and autophagosome membranes. Mammalian cells have at least six ATG8s, classified into LC3 and GABARAP families, while yeast only has one (8). It should be noted that ATG8 proteins are thought to be distributed on both the inner and outer membrane of the phagophore throughout autophagy process. Those localised on the inner surface of the phagophore recruit autophagy cargo receptors and get degraded in lysosomes. For example, p62, the first identified cargo receptor (9), in the cytosol can be degraded in response to amino acid starvation, a bulk autophagy induction signal (10). Alternatively, during damage or stress signals inducing selective autophagy, p62 binds polyubiquitin chains and forms droplet-like structures (6). In all forms, p62 interacts with ATG8 via a consensus motif, called the LIR (LC3-interacting region) motif. Upon deletion of the p62 LIR motif (7) or overexpression of the HyD-LIR probe (11), which has stronger binding affinity to LC3 compared to p62 (6), phagophore membranes are not able to enwrap p62 droplets, and lead to membrane bending away from the droplets.

A growing number of *in vitro* reconstitution studies have expanded our understanding of the molecular mechanisms of autophagy, from the initiation steps to cargo recruitment (reviewed in 12, 13). Giant unilamellar vesicles (GUVs) are commonly used and ideal for microscopic analysis to visualise the protein-membrane interaction and membrane deformation during ATG8 lipidation and cargo recruitment (examples in 14–18). Reconstitution of ATG8-PE conjugation is a fundamental and key approach, not only to determine the function of the upstream regulators, but also to study the downstream cargo recruitment.

While ATG8-PE conjugation has been reconstituted on the outer surface of GUV membranes, phagophore membranes are highly dynamic and have more complex topology (3, 5). There are three main configurations: the tips of phagophore with high positive membrane curvature, the outer membrane with positive membrane curvature decorated with ATG8-PE conjugates and recruited scaffold proteins, and the inner membrane with negative membrane curvature that has ATG8-PE conjugates with bound receptors and cargo. To gain insight into the functional consequences of the topologically different surfaces, the regulation of cargo recruitment and ATG8 distribution on the inside of the membrane, a model membrane that recapitulates these topologies is required.

Here we reconstituted ATG8 lipidation and ATG8-mediated cargo recruitment using a number of *in vitro* reconstitution approaches, including protein encapsulation within GUVs and a bead-based membrane expansion assay. Expanding on previous studies, we generated a membrane platform to encapsulate proteins of interest inside GUVs mimicking the inner leaflet of the phagophore membrane. We first reconstituted WIPI2b-directed ATG8 lipidation inside the GUVs and found a strict dependence of ATG8 lipidation on WIPI2b and the presence of PI3P. However, we found p62 can alternatively promote ATG8 lipidation in the absence of WIPI2b-PI3P interaction, consistent with previous studies showing that autophagy cargo may function upstream of ATG8 lipidation and direct autophagosome formation, reviewed in (19). Furthermore, focusing on the interactions between ATG8s and p62 on the membrane, we showed the interactions between ATG8 and p62 or p62 droplets were highly dynamic and mediated in a LIR-dependent manner. Interestingly, using the bead-based membrane expansion assay, we found that ATG8 coated GUV membranes underwent a deformation and wrapped around the p62 droplets bound to the beads. Additionally, ATG8 redistributed across the membrane and became locally enriched on the surface of the droplets. Our results reveal a key feature of inner phagophore membranes, where cargo receptors can concentrate lipidated ATG8, and thus shape and direct phagophore membrane expansion.

## Results

### Characterisation of WIPI2b-directed ATG8 conjugation machinery

The canonical ATG8 lipidation machinery is modulated in a PI3P-dependent manner by WIPI proteins (20–22). To better understand this pathway, we purified full-length human WIPI2b, an upstream regulator of ATG12-ATG5—ATG16L1 complex (hereafter, the E3 complex). To assess its lipid binding activity, we performed a liposome flotation assay using purified WIPI2b and liposomes (∼100nm in size) composed of various lipid compositions (**Fig. 1A-B**). Consistent with previous studies on WIPI proteins (21, 23), WIPI2b does not bind to liver PI but specifically binds to PI3P. Lipid packing defects caused by increasing the amount of DOPE (0%, 20% and 40%) in liposomes did not facilitate WIPI2b binding to the membrane. Additionally, we show that while the E3 complex alone did not bind to the liposomes containing 5% PI3P/20% DOPE/75% POPC, it was recruited to the liposome membrane with WIPI2b.

**Fig. 1.**
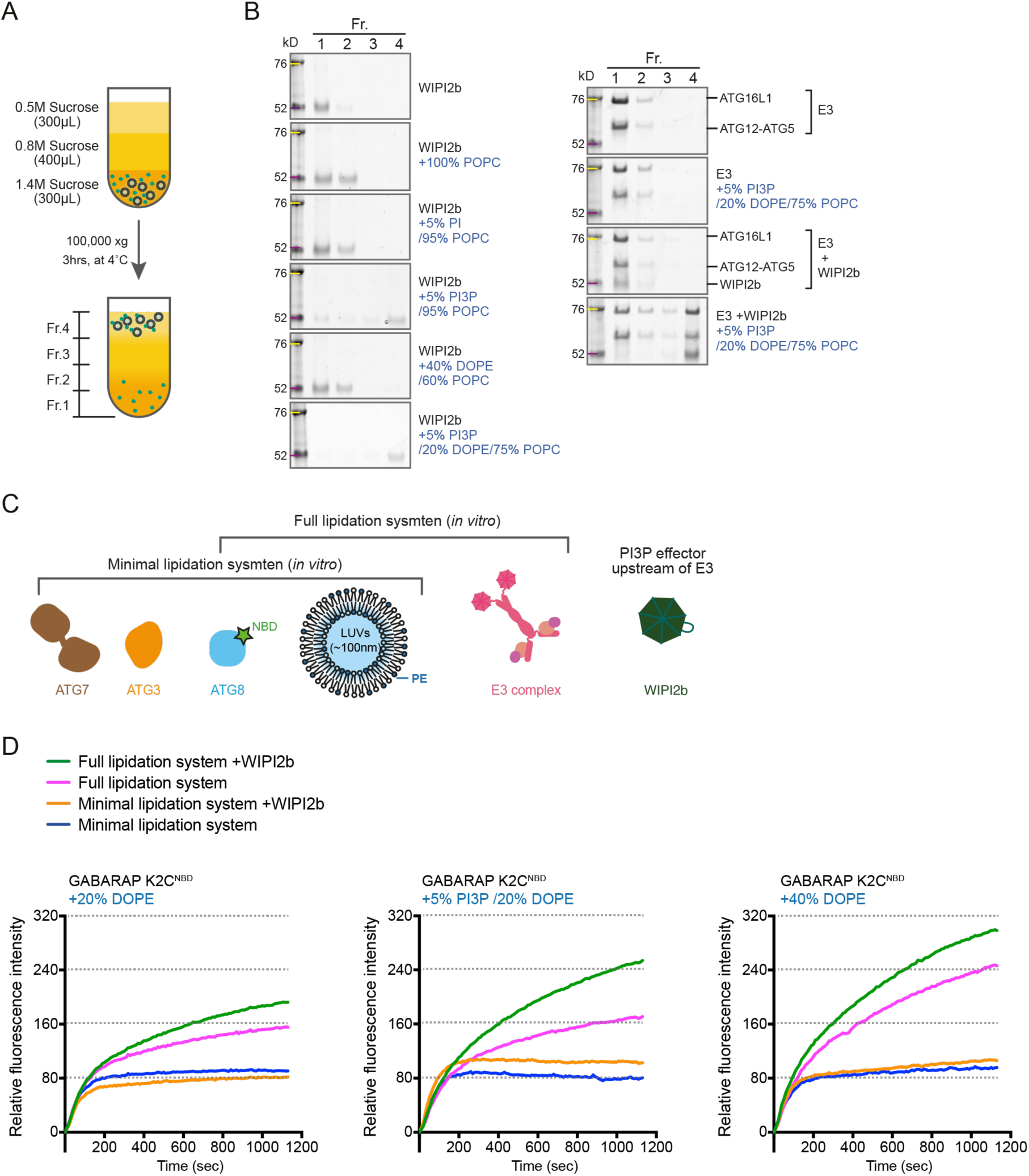
Characterisation of WIPI2b-directed ATG8-PE conjugation machinery. **(A)** Schematic diagram for flotation assay. **(B)** Liposome flotation assay of WIPI2b. Either 0.2μM WIPI2b, 0.2μM the E3 complex, or a mixture of 0.2μM WIPI2b and 0.2μM the E3 complex was mixed 0.5mM liposomes containing the indicated lipid composition. The protein-liposome mixtures were combined with 1.4M sucrose and overlaid with a sucrose gradient. After centrifugation, four fractions were collected from bottom to top and analysed by SDS-PAGE**. (C)** Schematic diagram for the minimal and full catalysts for ATG8 lipidation reaction. The activity of WIPI2b is assessed by the real-time lipidation assay, using N-terminal NBD-labelled ATG8, GABARAP K2C^NBD^. **(D)** NBD fluorescence changes of GABARAP K2C^NBD^ during the lipidation assay. “Full lipidation system + WIPI2b” contains 0.2μM ATG7, 0.2μM ATG3, 1μM NBD-labelled GABARAP, 0.05μM the E3 complex, 0.1μM WIPI2b and 1 mM liposomes. The other conditions were performed with the components as indicated in **(B).** The reaction was initiated by adding 1mM ATP/MgCl_2_ (time point at 0s). The relative fluorescence intensity was normalised to “Full lipidation system + WIPI2b” condition without ATP. Data represent mean values (n = 3).

To further assess WIPI2b activity in the ATG8 lipidation reaction, we employed a real-time *in vitro* assay by using GABARAP labelled with 7-nitrobenz-2-oxa-1,3-diazol-4-yl (NBD) at its N-terminal residue: GABARAP K2C^NBD^ (**Fig. 1C and Fig. S1A;** (24)). *In vitro*, the minimal catalysts for ATG8-PE conjugation are ATG7 and ATG3 (‘Minimal lipidation system’) (25, 26), whereas the addition of the E3 complex (‘Full lipidation system’) facilitates more efficient ATG8-PE conjugation (21, 24, 27). We employed liposomes with the following compositions: 20% DOPE/80% POPC to mimic PE levels in the ER (28); 20% DOPE/5% PI3P/75% POPC to mimic the phagophore membrane after PI3P generation by the PI3K complex (21, 29); 40% DOPE/60% POPC, which is a lipidation-prone condition due to its high PE content (30) (**Fig.1D and Fig. S1B**). We found that regardless of the lipid compositions, the addition of WIPI2b to the full lipidation system increased the NBD signal compared to the full lipidation system alone, suggesting that WIPI2b activated the E3 complex. With the same amount of the PE, the presence of PI3P led to an even higher NBD signal when WIPI2b was added to the full lipidation system, supporting its role in both activating and recruiting the E3 complex to the membrane (**Fig. 1D and Fig. S1C**). Taken together, our *in vitro* characterisation of WIPI2b aligns with its known cellular functions and supports the recent findings that it allosterically activates the E3 complex to promote ATG8 lipidation (20, 21, 27).

### Reconstitution of WIPI2b-directed ATG8 conjugation machinery inside GUVs

Since autophagy cargo recruitment and sequestration occur on the inner leaflet of the phagophore membrane, and finally within an enclosed environment, the autophagosome, we employed a modified cDICE (continuous droplet interface crossing encapsulation) method (31, 32) to investigate this process inside a GUV. We first reconstituted the ATG8 lipidation reaction inside the GUVs to decorate the inner membrane surface with lipidated ATG8 (**Fig. 2A**). Purified ATG7, ATG3, and Alexa Fluor™ 555-labeled GABARAP K2C were incubated with or without ATP for 1hr before adding the E3 complex and WIPI2b (**Fig. 2B**; see Materials and Methods). This reaction mixture was then injected into the oil phase containing different lipid compositions: 20% POPE/5% PI3P/74.5% POPC/0.5% NBD-PE or 20% POPE/79.5% POPC/0.5% NBD-PE. Maintaining the same amount of PE, the addition of PI3P allowed us to assess the WIPI2b activity inside the GUVs.

**Fig. 2.**
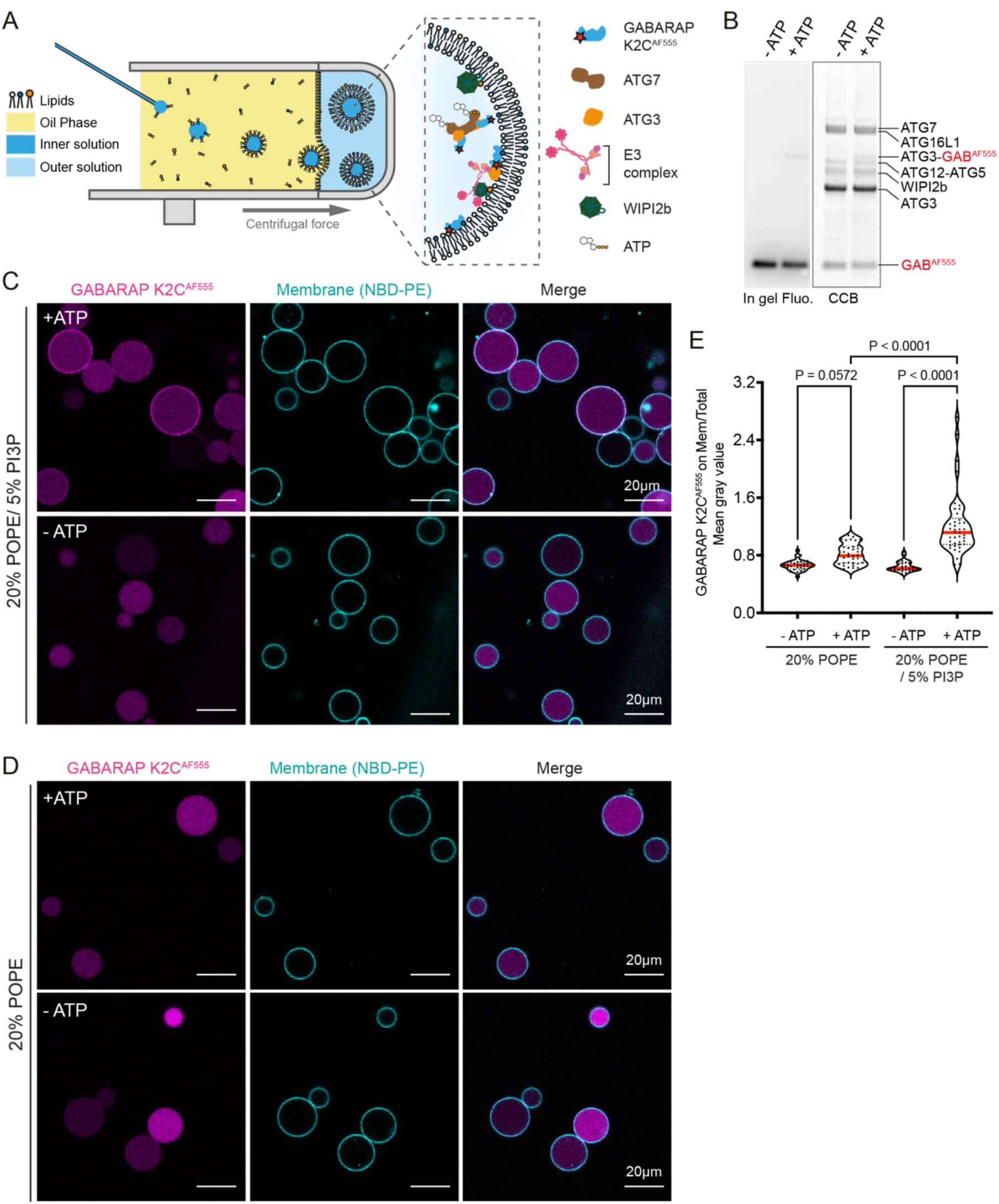
Reconstitution of WIPI2b-directed ATG8-PE conjugation reaction within GUVs. **(A)** Schematic diagram for cDICE. The inner solution containing proteins of interest is injected into a spinning chamber via a glass capillary. The chamber was successively filled with aqueous outer solution and lipid-in-oil mixture. The inner leaflet of GUV is formed when the inner solution droplet is flying through the oil phase and the outer leaflet is formed when the droplet passes through the oil-water interface. **(B)** Full lipidation system with WIPI2b, with or without ATP/MgCl_2_. The inner solution contained 0.5μM ATG7, 1.5μM ATG3, 0.5μM the E3 complex, 1μM WIPI2b, 0.5μM GABARAP K2C^AF555^, prepared with or without 1mM ATP/MgCl_2_. **(C)** Representative images show GABARAP-PE conjugation within the GUVs containing 20% POPE/5% PI3P/74.5% POPC/0.5% NBD-PE. **(D)** Representative images show GABARAP-PE conjugation within the GUVs containing 20% POPE/79.5% POPC/0.5% NBD-PE. **(E)** Quantification of GABARAP fluorescence on the membrane. The fluorescent intensity of GABARAP on the membrane compared to that of total GABARAP encapsulated inside the GUVs (20% POPE: -ATP n=34, +ATP n=34; 20% POPE/5% PI3P: -ATP n=27, + ATP n=42). The thick red lines and doted black lines in the violin plot represents the medians and quartiles, respectively.

Following the successful encapsulation of the WIPI2b-directed ATG8 lipidation machinery inside the GUVs, we found that GABARAP was conjugated only to the GUV membranes containing PI3P (**Fig. 2C-D**). The conjugation reaction inside the GUV membranes was facilitated by WIPI2b-PI3P binding, which recruited the E3 complex and promoted ATG8-PE conjugation (**Fig. S2A-B**). Extending the results from previous ATG8 reconstitution studies on the outer surface of liposomes (15, 21, 22), the cDICE encapsulation system, for the first time, reconstituted ATG8 lipidation on the inner leaflet within the confined environment of a GUV. Furthermore, our results also demonstrated the strict dependence on WIPI2b-PI3P binding, which is crucial for an effective ATG8 lipidation reaction.

### Reconstitution of p62-directed ATG8 lipidation reaction inside GUVs

We next investigated the effect of the autophagic cargo receptor p62 on the ATG8 lipidation machinery reconstituted within the GUVs. We prepared the WIPI2b-directed ATG8 conjugation reaction mix with or without addition of purified EGFP-p62. As described above, we found the ATG8 lipidation strictly depends on WIPI2b-PI3P binding (**Fig. S3A and S3C)**. Inside the GUVs containing 20% POPE and 5% PI3P, the presence of p62 did not affect GABARAP recruitment to the membrane. However, within the GUVs containing 20% POPE but no PI3P, p62 was recruited to the membrane and facilitated GABARAP lipidation in a manner independent of WIPI2b-PI3P (**Fig. S3B and C**). In addition, p62 was recruited to the membrane by lipidated GABARAP (**Fig. S3B**).

### Recruitment of p62 and p62 droplets to ATG8s on the inner membrane depends on its LIR motif

Having successfully reconstituted ATG8 lipidation and p62 recruitment inside the GUVs, we sought to use this system to explore ATG8 and more relevant cargo interactions. It was previously reported that cargo receptors trigger ATG8 lipidation via interaction with the E3 complex, WIPI proteins and/or other upstream regulators of ATG8 lipidation machinery (15, 33, 34). Thus, to focus on the downstream interaction between ATG8-cargo on the inner leaflet of the membrane, we employed a simplified system using C-terminal His_6_-tagged ATG8 and GUVs containing nickel lipids to anchor the ATG8 on the inner membrane and mimic the lipidated status (**Fig. 3A**).

**Fig. 3.**
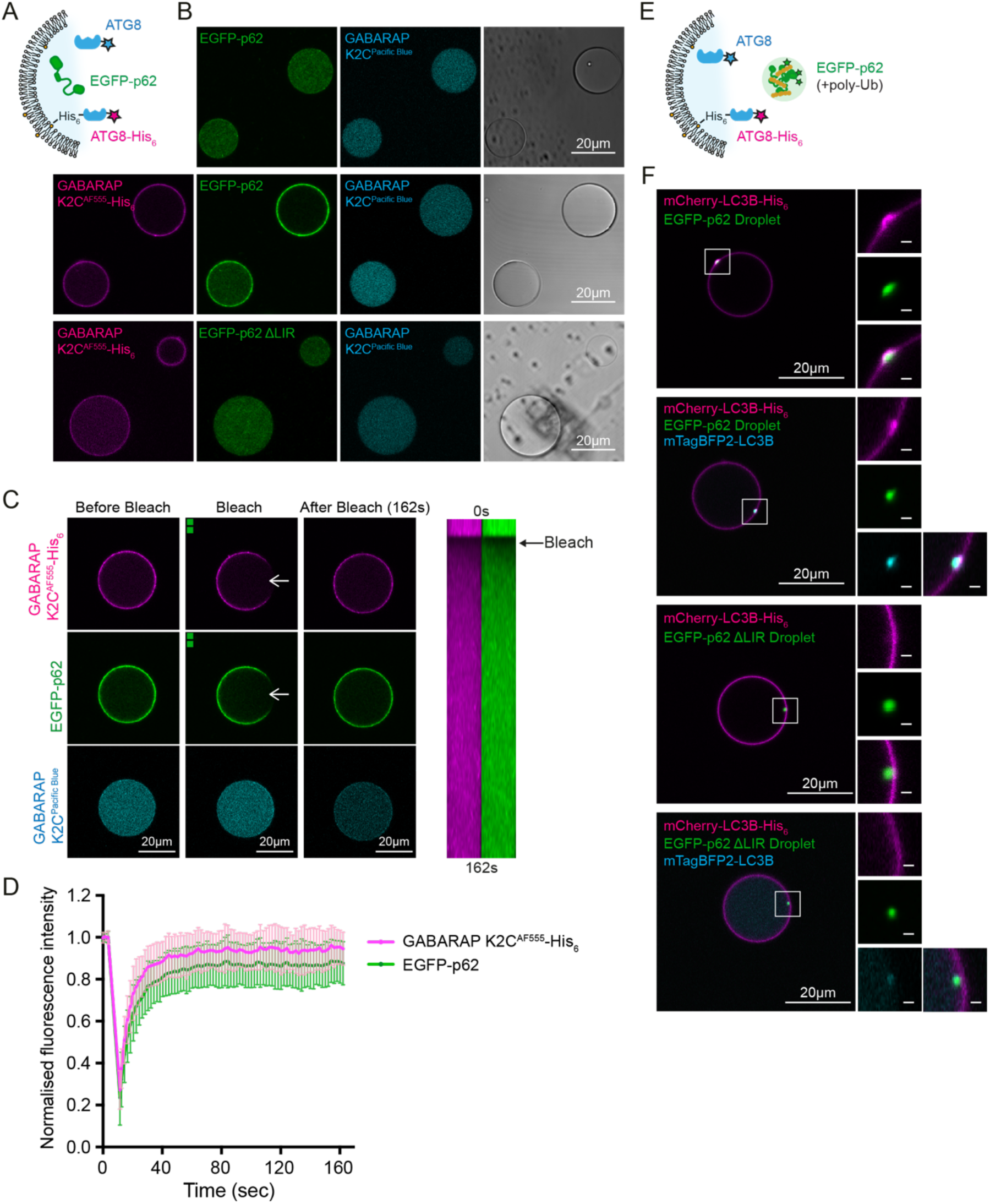
Soluble and droplet-like p62 is recruited to membrane depending on its LIR motif. **(A)** A simplified system used to investigate p62 recruitment to membrane-bound ATG8 and soluble ATG8. **(B)** GUVs composed of 95% POPC and 5% DGS Ni-NTA encapsulated 5μM EGFP-p62 WT or ΔLIR with 1μM GABARAP K2C^Pacific Blue^ and/or 1μM GABARAP K2C^AF555^-His_6_. **(C)** FRAP of membrane bound GABARAP and EGFP-p62. Kymographs shows the fluorescence recovery in the photobleached area, indicated by the white arrows. **(D)** Quantification of fluorescence intensity recovery of GABARAP K2C^AF555^-His_6_ and EGFP-p62 (mean ± SD, n=25 GUVs). **(E)** Reconstitution of p62 droplet recruitment to ATG8 decorated membrane. **(F)** EGFP-p62 WT or ΔLIR droplets were encapsulated inside the GUVs with 1μM mCherry-LC3B-His_6_ and/or 1μM mTagBFP2-LC3B. GUV membranes contained 95% POPC and 5% DGS Ni-NTA. Magnified views of the outlined regions are shown in the left panels. Scale bars: 2μm

Encapsulated GABARAP K2C^Pacific Blue^ and EGFP-p62 remained diffuse within the GUV with no membrane interaction, while GABARAP K2C^AF555^-His_6_ which bound the nickel lipids on the membrane, recruited EGFP-p62 to the membrane (**Fig. 3B**). Furthermore, deletion of the LIR motif in EGFP-p62 impaired p62 recruitment to the membrane, showing as expected that p62 recruitment to ATG8-decorated membrane depends on its LIR motif. We also assessed the dynamics of ATG8-p62 interaction on the membrane using fluorescence recovery after photobleaching (FRAP) assay. With a finite amount of the proteins in a confined environment, we found that fluorescently labelled GABARAP K2C^AF555^-His_6_ and EGFP-p62 were distributed uniformly across the membrane. Both proteins recovered their fluorescence with a similar kinetics after photobleaching (**Fig. 3C-D**), indicating that ATG8 and p62 on the membrane are highly mobile.

We then asked if p62 condensates formed with K63-linked ubiquitin chains could bind ATG8s in our encapsulated system. p62 specifically binds to K63-linked polyubiquitin chains and undergoes liquid-liquid phase separation and forms biomolecular condensates in cells and *in vitro* (35, 36). First, we reconstituted p62 phase separation induced by K63-linked polyubiquitin chains (**Fig. S4**; see Materials and Methods) to form p62 “droplets” (36) equivalent to p62 bodies in cells (37, 38). EGFP-p62 underwent phase separation immediately upon mixing with K63-linked polyubiquitin chains, which was promoted by addition of 2% PEG8K (**Fig. S4A-C**). In line with previous studies (36, 39), we showed that p62 droplets sequester soluble GABARAP K2C^AF555^ in a LIR-dependent manner (**Fig. S4D**).

Next, we investigated the interaction of the p62 droplets with both membrane-bound and soluble ATG8 (**Fig. 3E**). Here, we used fluorescently tagged LC3B, and although it has a weaker binding affinity to p62 compared to GABARAP (40), it has been widely used for *in vitro* reconstitution studies of autophagy (15, 21). We found that p62 droplets were recruited by membrane-bound mCherry-LC3B-His_6_ (**Fig.3F**). Interestingly, mCherry-LC3B-His_6_ was distributed across the membrane but appeared to be locally enriched on the surface of the p62 droplets. A small membrane deformation of the LC3-bound p62 droplet positive GUV membrane was reproducibly detected (**Fig. 3F**). To mimic the conditions in cells, where both soluble and membrane-bound LC3B are present, we then encapsulated both mCherry-LC3B-His_6_ and soluble mTagBFP2-LC3B within the GUV. We found p62 droplets were recruited to the membrane, and also concentrated the membrane-bound LC3B on the surface of the droplet as well as the soluble LC3B within the droplet (**Fig.3F**). Notably, during p62 droplet formation, p62 was highly concentrated in a dense structure, which is capable of capturing and concentrating soluble ATG8 inside the droplets (**Fig. S4D**). Deletion of p62 LIR motif abolished its interaction with both membrane-bound LC3B and soluble LC3B. Taken together, our results reveal a highly dynamic interaction and distinct behaviours, between soluble and membrane bound forms of ATG8 with p62 and p62 bodies.

### Interaction between p62 and membrane-bound ATG8 directs membrane expansion

We successfully encapsulated p62 droplets inside the GUVs, reconstituted cargo recruitment on the inner leaflet of the membranes and showed that binding of p62 droplets to membrane bound LC3B deforms the membrane. We next developed a second approach, a novel bead-based membrane expansion assay to confirm and further investigate how this interaction directs membrane deformation (**Fig. 4 and Fig. S5**). Briefly, we coated the GFP-TRAP beads with soluble EGFP-p62 or EGFP-p62 droplets and then added GUV membranes prepared with nickel lipids and decorated with mCherry-LC3B-His_6_ (**Fig. 4A and C**). When the beads were coated with soluble p62, the LC3B-GUV membranes bound to the beads. On the GUV, the LC3B was redistributed and locally enriched at the interaction interface with p62, and the membrane was deformed at the contact site (**Fig. 4B**). Interestingly, in the case of p62 droplet-bound beads, membrane-bead binding occurred, and on the GUVs, we observed additional interactions: the GUV-LC3B membrane was reshaped and wrapped around the p62 droplets. In line with the results shown in **Fig. 3F**, LC3B was redistributed and was enriched at the surface of the p62 droplets (**Fig.4D**). The membrane deformation and ATG8 redistribution on the membrane was mediated by the p62 LIR motif (**Fig. 4E and F**). Deletion of p62 LIR motif impaired the interaction between membrane-bound LC3B with soluble p62 and as well as with p62 droplets. Importantly, no membrane bending was observed without the LIR motif. In summary, using a cargo template with uniform (soluble) p62 or polyubiquitin-induced p62 droplets, we recapitulated membrane deformation and ATG8 redistribution across the membranes, mediated by ATG8 interactions with the cargo receptor. Importantly, our *in vitro* assays can be used to dissect these interactions that occur in cells during phagophore membrane expansion and cargo recruitment.

**Fig. 4.**
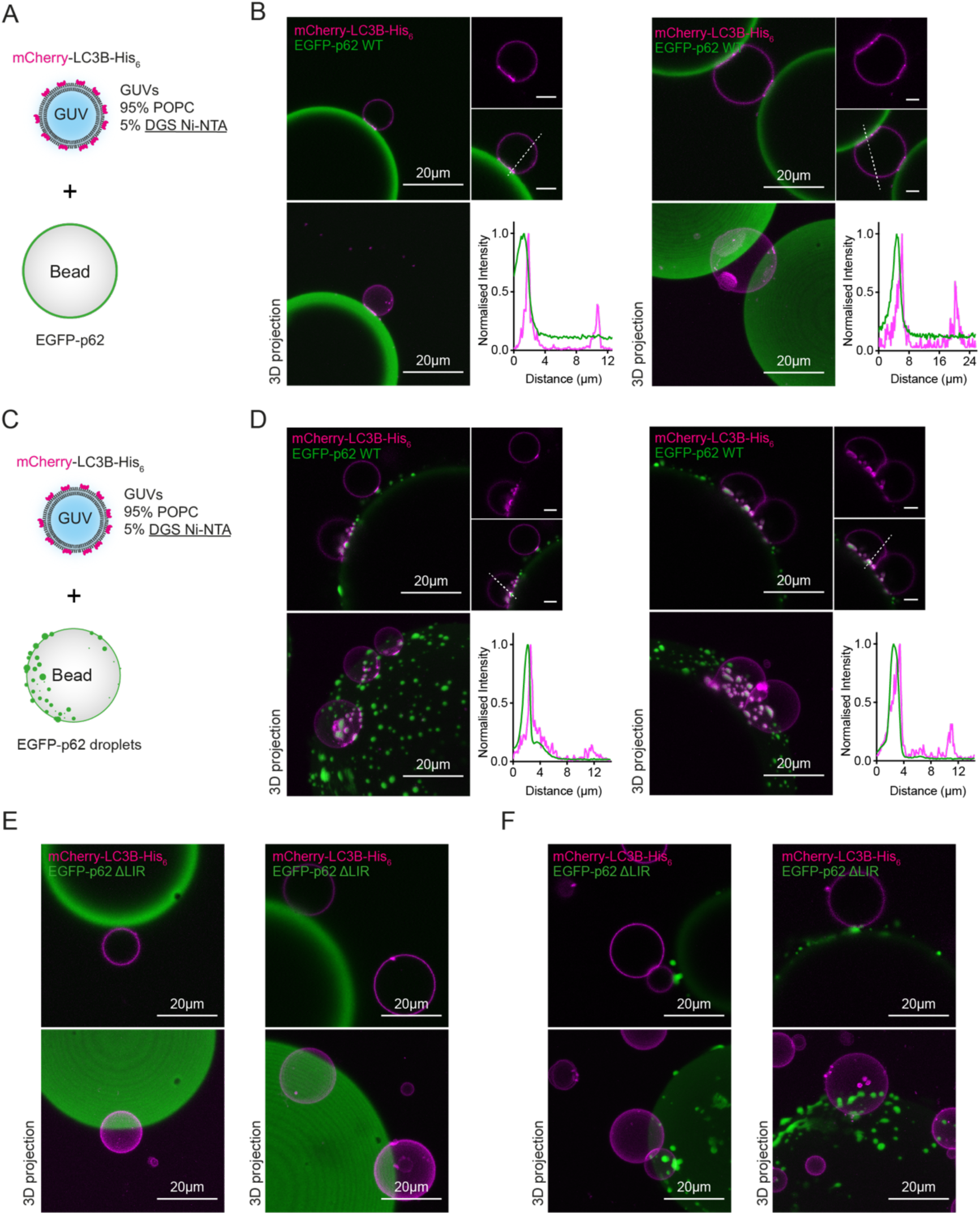
p62-ATG8 interaction directs membrane bending. **(A) and (C)** Experimental design of the bead-based membrane expansion assay. Soluble or droplet-like EGFP-p62 was coated on GFP-TRAP beads and mixed with GUVs (95% POPC/ 5% DGS Ni-NTA) decorated with mCherry-LC3B-His_6_. **(B) and (D**) Confocal images showing membrane bending directed by the interaction between mCherry-LC3B-His_6_ and p62 WT or p62 WT droplets on the bead. Magnified views of the GUV regions are shown in the left panel. Scale bars, 5μm. 3D projections of the GUV-bead interaction regions are shown at the bottom. Intensity profile shows the distribution of mCherry-LC3B-His_6_ across the membrane towards the contact sites with p62 or p62 droplets on the beads. **(E) and (F)** Confocal images showing random attachment of the GUVs decorated with mCherry-LC3B-His_6_ to p62 ΔLIR or p62 ΔLIR droplets on beads. 3D projections of the GUV-bead interaction regions are shown at the bottom.

## Discussion

Recently, several labs have made use of ATG8-PE reconstitution assays on the outer surface of GUVs as a readout to assess the function of upstream regulatory autophagy machinery (15, 21), or as membrane platforms to investigate downstream cargo recruitment (33, 41). In this study, we have successfully reconstituted WIPI2b-directed ATG8 lipidation and ATG8-mediated cargo recruitment inside GUVs to mimic the inner membrane of a growing phagophore. In addition, we could reconstitute and test the effect of two p62-based cargoes, p62 and polyubiquitin-induced p62 droplet, on the distribution of the membrane bound ATG8s.

Having demonstrated the crucial function of WIPI2b for initiating autophagy cascade (**Fig. 1**) and using a real-time lipidation assay with ∼100nm liposomes, we showed that WIPI2b facilitates ATG8 lipidation and dissected its role in directing the E3 complex to the PI3P-containing membranes and activating the E3 complex allosterically.

Using the cDICE approach, we reconstituted WIPI2b-directed (**Fig.2**) and cargo receptor p62-directed (**Fig. S3**) ATG8 lipidation. This experimental system based on encapsulation within a GUV provides an alternative approach to the use of ∼100nm liposomes which present a highly positive membrane curvature, as the GUV membranes are almost flat. Previous reconstitution studies of ATG8 lipidation on GUVs have either required a high percentage of PE (≥30%) with negative charged lipids, such as PI or PS in the absence of PI3P and WIPI2 (16–18), or PE levels similar to the ER (∼20-25%) with PI3P and WIPI2 (21). Upon encapsulation of the reaction mixture containing the conjugation machinery, the E1-E2-E3 and WIPI2b within the GUVs, the simple lipid compositions (20% POPE vs. 20% POPE/5% PI3P, supplemented with POPC), supported ATG8 lipidation which was strictly dependent on the interaction between WIPI2b and PI3P (**Fig.2 and Fig. S2**). Furthermore, addition of p62 to the E1-E2-E3-WIPI2b reaction within the GUVs facilitated ATG8 lipidation on the membrane containing 20% POPE without 5% PI3P (**Fig.S3**). These results demonstrated a cargo-directed ATG8 lipidation mechanism independent of WIPI2b-PI3P interaction. This mechanism could be explained by the potential of purified p62 to spontaneously form high-order oligomers, generating clusters of LIR motifs that interact with multiple ATG8s, in support of the previous data that p62 oligomers locally concentrate the ATG8s and promote ATG8 lipidation (42, 43). Since the amount of proteins injected into the GUVs was finite, the reaction rate of ATG8-PE conjugation on GUV membranes containing 20% POPE/5% PI3P may reach a plateau under our reaction conditions independent of p62 oligomerisation.

We further focused on autophagy cargo recruitment inside the GUVs with a simplified system by generating GUV vesicles with nickel-chelating lipids and encapsulating His_6_-tagged fluorescent ATG8 proteins. We succeeded in visualising the dynamic diffusion of membrane-bound ATG8 and p62 (**Fig.3**). Both soluble p62 and p62 droplets were recruited to ATG8-bound membranes via their LIR motif. When encapsulating p62 droplets inside the GUVs, we observed a local enrichment of membrane-bound ATG8 on the surface of p62 droplets and soluble ATG8 sequestered within the p62 droplets (**Fig. 3F**). Due to the complexity of cDICE approaches, we did not manage to encapsulate the polyubiquitin synthesis reaction, p62 and the WIPI2b-directed ATG8 lipidation machinery inside the GUVs to examine how formation of p62 droplets mediate ATG8 lipidation. However, based on our current results and previous studies (34, 36, 39, 44), it is reasonable to hypothesise that the p62 droplets may concentrate the ATG8 conjugation machinery and promote ATG8 lipidation at the contacts between the droplets and the membranes.

In addition, we have developed a bead-based membrane expansion assay (**Fig. 4 and Fig. S5**). Our results indicated that the ATG8 is dynamically redistributed across the outer membrane of GUVs upon binding to p62 and most strikingly to p62 droplets. These results suggest and support a reorganisation and local enrichment of lipidated ATG8 on the concave site of the phagophore membrane, where cargo recruitment occurs. Further development and optimisation is required, but our assays also provide promising tools to investigate various types of selective autophagy by generating an uneven distributed cargo templet and mimicking local enrichment of ubiquitin signals during autophagy process (45).

## Materials and Methods

### Materials

Plasmids used in this study are provided in <u>SI Appendix, Table S1</u>. All lipids were obtained from Avanti Polar Lipids: POPC (850457C), DOPE(850725C), POPE (850757C), PI3P (850150P), Liver-PI (840042C), DGS-Ni-NTA (790404C), 18:1 Liss Rhod PE (810150C), 18:1 NBD PE (810145C). Marina Blue^TM^ DHPE (M12652) was obtained from Invitrogen. For giant unilamellar vesicle preparation, silicon oil (317667), mineral oil (M5904) and anhydrous chloroform (372978) were obtained from Sigma-Aldrich. For protein labelling, IANBD (D2004), Pacific Blue™ C_5_-Maleimide (P30506), Alexa Fluor™ 488 C_5_-Maleimide (A10254), and Alexa Fluor™ 555 C_2_-Maleimide (A20346) were obtained from Invitrogen. Antibodies used for immunoblotting: rabbit monoclonal anti-GABARAP (Cell Signaling Technology, 13733), and HRP-conjugated anti-rabbit IgG (GE Healthcare, #NA934). ATP solution (100 mM, R0441) was purchased from Thermo Fisher. Bovine ubiquitin (U6253) was purchased from Sigma-Aldrich.

### Recombinant protein expression and purification

Plasmids for bacterial expression were transformed in *Escherichia coli* BL21 (DE3) cells. Bacteria were grown in LB medium at 37℃ until an OD_600_ of 0.6 and protein expression was induced with 0.75mM IPTG for 16hrs at 18℃. The GST-tagged proteins were purified as described before (24, 40). Briefly, cells were harvested and lysed in 50mM Tris-HCl pH8.0, 500mM NaCl, 0.5mM TCEP, 0.4mM AEBSF, 15μg/ml benzamidine and 0.1% TritonX-100, followed by tip sonication. Lysates were then cleared by centrifugation at 25,000 × g for 30 min at 4°C. The GST-tagged proteins were extracted with Glutathione-Sepharose 4B affinity matrix (GE Healthcare) for 1.5-2 hr and recovered by 3C protease cleavage at 4°C overnight in 50mM Tris-HCl pH 8.0, 500mM NaCl, and 0.5mM TCEP. His_6_-mUBE1 was purified as previously described (46) with minor modification. The expressed protein was absorbed by HisPur^TM^ Cobalt Resin for 2 hr incubation at 4°C and then eluted in 50mM Tris-HCl, pH8.0, 500mM NaCl, 0.5mM TCEP and 300mM imidazole. All the proteins were further purified by size-exclusion chromatography using Superdex 200 16/60 column (GE Healthcare) equilibrated in buffer containing 25mM Tris-HCl pH8.0, 150mM NaCl, and 0.5mM TCEP.

ATG7 and WIPI2b were expressed and purified from Sf9 cells as previously described (24). Briefly, pBacPAK-His_3_-GST-ATG7 or pBacPAK-His_3_-GST-WIPI2b were transfected into insect cells Sf9 using flashBAC GOLD (Oxford Expression Technologies [OET]) and Fugene HD transfection reagent (Promega) according to the manufacturer’s instructions. For protein expression, Sf9 cells were infected with the baculovirus harbouring His_3_-GST-ATG7 or His_3_-GST-WIPI2b in Sf-900 II SFM medium (Thermo). To purify ATG7, the cells were resuspended in buffer containing 50mM Tris-HCl pH 8.0, 500mM NaCl, 0.5mM TCEP and EDTA-free cOmplete Protease Inhibitor cocktail (Roche) and lysed by tip sonication. The remaining purification steps followed the same procedure as described above. When purifying WIPI2b, the cells were lysed in buffer containing 50mM Tris-HCl pH 7.5, 500mM NaCl, 2mM MgCl_2_, 0.5mM TCEP, 10% glycerol, 0.5% Triton and EDTA-free cOmplete Protease Inhibitor cocktail (Roche). GST-tagged WIPI2b was absorbed by Glutathione-Sepharose 4B affinity matrix (GE Healthcare) for 1.5 hr and GST was excised by 3C protease at 4°C overnight in 50mM Tris-HCl pH 7.5, 500mM NaCl, 2mM MgCl_2_, 0.5mM TCEP, 10% glycerol. The eluted protein was then purified with the Superdex 200 10/300 GL column equilibrated in buffer containing 25mM Tris-HCl pH 7.5, 150mM NaCl, 1mM MgCl_2_, 0.5mM TCEP, 10% glycerol.

Human ATG12-ATG5—ATG16L1β (the E3 complex) was either purified from High Five insect cells as previously described (24).

### Protein fluorescent labelling

Pacific Blue™ C_5_-Maleimide or Alexa Fluor™ 555 C_2_-Maleimide were conjugated to the cysteine residue introduced at GABARAP N-terminal lysine 2 (GABARAP K2C). Alexa Fluor™ 488 C_5_-Maleimide was conjugated to the native cysteine residues in WIPI2b. The purified proteins were transferred to labelling buffer (20mM Tris-HCl 7.5 and 150mM NaCl) using Zeba™ Spin Desalting Columns (Thermo), immediately prior to labelling. The proteins were then labelled with an equimolar amount of fluorescent dyes for 1hr at room temperature in the dark. Unlabelled dyes were removed by Zeba™ Spin Desalting Columns (Thermo), preequilibrated with 25 mM Tris-HCl pH 8.0, 150 mM NaCl, and 0.5 mM TCEP. The concentration of labelled proteins and labelling efficiency were estimated by NanoDrop Spectrophotometer. NBD labelling of GABARAP K2C was performed as previously described (24). Fluorescent labelled proteins were snap-frozen in liquid nitrogen and stored at −80℃.

### Liposome preparations

Lipids with desired molar ratio were mix in chloroform, dried under N_2_ gas, and further desiccated for 2hrs under vacuum. The lipid film was resuspended by vortexing in the assay buffer (25mM Tris-HCl pH 8.0, 150mM NaCl, and 0.5mM TCEP) and subjected to five times freeze-thaw cycles in liquid nitrogen and water batch. The resuspended lipids then were extruded 21 times through 0.2μm membrane followed by at least 41 times through 0.1μm membrane (Whatman) using a Mini-Extruder (Avanti Polar Lipid). The size of liposomes was measured by Zetasizer Nano ZS (Malvern Instruments). The liposomes had an average diameter of 100 nm. The final concentration of liposomes was 2mM.

### Flotation assay

300μL of bottom mixture was prepared at room temperature containing either 0.2μM WIPI2b, 0.2μM the E3 complex, or 0.2μM WIPI2b and 0.2μM the E3 complex, along with 0.5mM liposomes and 1.4M sucrose in assay buffer (25 mM Tris-HCl pH 8.0, 150 mM NaCl, and 0.5 mM TCEP). The bottom mixture was immediately transferred into centrifugation tube (Beckman Coulter, #343778), overlaid with 400μL of 0.8M sucrose in assay buffer and topped with 300μL of 0.5M sucrose in assay buffer. Gradients were centrifugated at 100,000 x g (47,000 r.p.m.) in a TLA120.2 rotor at 4℃ for 3hrs. Four fractions (each fraction ∼250μL) were recovered, starting from the bottom. Each fraction was incubated with 10μL StrataClean resin (Agilent) for 30min at 4℃ to concentrate the diluted proteins in each fraction. The resin was pelleted and mixed with 10μL SDS-PAGE sample buffer, heated at 95 for 5min, and then subjected to SDS-PAGE followed by Coomassie blue staining.

### Real-time lipidation assays

The real-time lipidation assays were performed as previously described (24). The NBD fluorescence (ex/em 468nm/535nm) was recorded using FP-8300 spectrofluorometer (JASCO). The excitation bandwidth was set to 5 nm, and the emission bandwidth was set to 10 nm. The reaction mix (80μL) contained 0.2μM ATG7, 0.2μM ATG3, 1μM NBD-labelled GABARAP, 0.05μM the E3 complex, 0.1μM WIPI2b and 1 mM liposomes. The reaction was initiated by addition of 1mM ATP/MgCl_2_. The control group contained all the proteins and liposomes, except ATP/MgCl_2_. The fluorescence increase (ΔEm535 nm) at each time point was calculated by subtracting the NBD signal recorded from the control group.

By the end of the reaction, 40μL of reaction mix was taken immediately, mixed with 10μL SDS-PAGE sample buffer, and heated at 95℃ for 5min. 10μL of reaction mix was resolved by NuPAGE Bis-Tris 4–12% gels (Life Technologies) followed by immunoblotting. Additionally, 25μL of reaction mix was resolved by NuPAGE Bis-Tris 4–12% gels (Life Technologies) and subjected to SYPRO™ Ruby Protein Gel Stain (Invitrogen™).

### cDICE system and GUV generation

GUVs were produced using the cDICE methods as described (31, 32, 47, 48), with a modified protocol. A rotating chamber was 3D printed with Clear resin on Formlabs Form 2. A magnetic IKA magnetic stirrer served as a motor, after removing the heating plate to expose the motor shaft.

Lipids were mixed at the desired molar ratio in chloroform in a 20mL glass vial, dried under nitrogen gas and further vacuumed for 10min. Lipid film was then re-dissolved in 600μL chloroform. While mixing on a vortex mixer, 9.4mL of silicon oil/mineral oil mix (ratio 4:1) was slowly added to the lipid solution in the glass vial. Then, the lipid-in-oil mixture was vortexed thoroughly and subsequently sonicated in bath sonicator for 20min (Fisherbrand^TM^ P-Series, 37Hz, 100% power, Degas Mode). The final lipid concentration in the lipid-in-oil mixture was around 250μM.

To encapsulate purified proteins or reaction mixtures, proteins of interest were prepared in the inner solution, while empty GUVs were generated using only the inner solution. The inner solution was composed of 25mM Tris-HCl pH 8.0, 150mM NaCl, 2mM MgCl_2_, 0.5mM TCEP and 9% iodixanol (OptiPrep^TM^, Sigma Aldrich). The outer solution contained the assay buffer (25mM Tris-HCl pH 8.0, 150mM NaCl, 2mM MgCl_2_ and 0.5mM TCEP), and glucose solution (∼2M in Milli-Q H_2_O) was used to match the osmolality of the inner solution. The osmolality of the inner and outer solution were measured by Gonotec® Osmomat™ 3000. For successful GUV generation, the osmolality of the outer solution should be higher than that of the inner solution by 15-25mOsm/kg.

The inner solution was loaded into BD Luer-Lock™ 1-mL syringe and connected through tubing to a fused silica capillary (inner diameter: 100μm). The syringe was placed on a Harvard Apparatus syringe pump. 600-800μL of outer solution was added into the rotating chamber, followed by the addition of the lipid-in-oil mixture to fill the chamber. The injection capillary was immersed into the oil phase and the inner solution containing the proteins of interest was injected at a flow rate of 150μL/hr for 10-15min. The GUVs were pipetted into a glass bottom microtiter plate (96-well or 384-well Greiner Bio-One), precoated with 5mg/mL BSA. For **Fig.2C** and **Fig. S2B**, GUVs were harvested and imaged in 96-well plates, while GUVs prepared for the other experiments were harvested and imaged in 384-well plates.

### *In vitro* lipidation of ATG8 proteins inside the GUVs

For **Fig.2C**, GABARAP K2C^AF555^ was preincubated ATG7, ATG3 and ATP/MgCl_2_ at 37℃ for 1hr in a volume of 100μL as previous described (18), before adding the E3 complex and WIPI2b. The reaction mixture was then diluted with the inner solution to 300μL to reach final concentrations of 0.5μM ATG7, 1.5μM ATG3, 0.5μM the E3 complex, 1μM WIPI2b, 0.5μM GABARAP K2C^AF555^ and 1mM ATP/MgCl_2_. 20μL of the final reaction mixture was checked by SDS-PAGE (**Fig. 2B**). The rest was loaded on the cDICE system.

For **Fig. S2B**, GABARAP K2C^Pacific Blue^ was preincubated ATG7, ATG3 and ATP/MgCl_2_ at 37℃ for 1hr as previous above in a volume of 100μL, before adding the E3 complex and WIPI2b. The reaction mixture was then diluted with the inner solution to 300μL to reach final concentrations of 0.5μM ATG7, 1.5μM ATG3, 0.5μM the E3 complex, 1.5μM WIPI2b (unlabelled to labelled at a molar ratio of 2:1), 0.5μM GABARAP K2C^Pacific Blue^ and 1mM ATP/MgCl_2_. 20μL of the final reaction mixture was checked by SDS-PAGE (**Fig. S2A**). The rest was loaded on the cDICE system.

For **Fig. S3**, GABARAP K2C^AF555^ was preincubated ATG7, ATG3 and ATP/MgCl_2_ at 37℃ for 1hr as described above in a volume of 100μL, before adding the E3 complex and WIPI2b. The reaction mixture was then diluted with the inner solution to 300μL to reach final concentrations of 0.5μM ATG7, 1.5μM ATG3, 0.5μM the E3 complex, 1μM WIPI2b, 0.5μM GABARAP K2C^AF555^, 1μM EGFP-p62 and 1mM ATP/MgCl_2_. The reaction mixture was then loaded on the cDICE system.

### *In vitro* K63-linked polyubiquitin chain synthesis and p62 droplet formation

The experiments were performed in the assay buffer (25mM Tris-HCl pH 8.0, 150mM NaCl, 0.5mM TCEP). To synthesise K63-linked polyubiquitin chains, the conjugation reaction containing 1μM His_6_-mUBE1, 8μM UBE2N, 8μM UBE2V1, 100μM ubiquitin was incubated in the presence of 5mM ATP/MgCl_2_ at 37℃ for 2hrs. The reaction mix was used immediately, or snap-frozen in liquid nitrogen and stored at −80℃ until the experiments.

For **Fig.S4C**, 5μM EGFP-p62 WT or ΔLIR was mixed with 20μL of the conjugation reaction, with or without 2% PEG8K in a reaction volume of 100μL. For **Fig. S4D**, 2.5μM EGFP-p62 WT or ΔLIR and 0.5μM GABARAP K2C^AF555^ was mixed with 30μL of the conjugation reaction with 2% PEG8K in a reaction volume of 100μL. The reaction mixtures was prepared and transferred immediately to 384-well glass bottomed plate (precoated with 5mg/mL BSA) and imaged by Zeiss LSM880 within 1hr.

### Soluble and droplet-like p62 recruitment assay

For **Fig. 3A**, 5μM EGFP-p62 WT or ΔLIR was mixed with 1μM GABARAP K2C^AF555^-His_6_ and 1μM GABARAP K2C^Pacific Blue^ in the inner solution. Then, soluble proteins were loaded on the cDICE system. For **Fig. 3C**, to encapsulate EGFP-p62 droplets inside the GUVs, the polyubiquitin conjugation reaction was prepared as described above in the inner solution. 5μM EGFP-p62 was mixed with 60μL of the polyubiquitin conjugation reaction, followed by the addition of 1μM mCherry-LC3B-His_6_ with or without 1μM mTagBFP2-LC3B, in a final inner solution volume of 300μL. The protein mixtures were then loaded on the cDICE system and used within 1hr.

### Bead-based membrane expansion assay

A mixture of 2.5μM EGFP-p62 (final concentration) and 60μL of the polyubiquitin reaction with or without ATP was incubated with 1.25μL of GFP-TRAP beads (ChromoTek) in assay buffer containing 25mM Tris-HCl pH 8.0, 150mM NaCl, 0.5mM TCEP, and 2% PEG8K, in a final volume of 250μL at 4℃ for 15min. The beads were washed three times with the outer solution containing 25mM Tris-HCl pH 8.0, 150mM NaCl, 2mM MgCl_2_, 0.5mM TCEP, and glucose solution (∼2M in Milli-Q H_2_O) adjusted to match the osmolality of the inner solution. After the final wash, the beads were resuspended into 100μL of the outer solution.

Empty GUVs (95%POPC/5% DGS Ni-NTA) were generated by the cDICE method as described above. 100μL of the GUV solution was collected, mixed with 100nM mCherry-LC3B-His_6_, and loaded into 384-well plate (precoated with 5mg/mL BSA). After incubation at room temperature for 20min, the GUVs were pelleted at the bottom of the plate and mCherry-LC3B-His_6_ was recruited to the membranes. Subsequently, 50μL of the supernatant was removed, and 50μL of the p62 or p62-droplet coated bead solution was added and gently mixed with the GUVs.

### Confocal microscopy

All the images were acquired using Zeiss LSM880, equipped with a 40x/1.2NA water objective at room temperature, with 512 x 512 pixel resolution. Random views were picked for imaging. Pacific Blue and mTagBFP2 were excited with 405nm laser. NBD-PE was excited with 456nm laser. Alexa Fluor 488 and EGFP were excited with 488nm laser. Alexa Fluor 555, mCherry and Rhod-PE were excited with 561nm laser. Z-stack images with 1μm intervals were processed using “Maximum Intensity Projection” function in Zeiss Zen software.

### Fluorescence recovery after photobleaching

FRAP experiments were performed with Zeiss LSM880. The images were acquired with a resolution of 512 x 512 pixels. Fluorescence of Alexa Fluor 555 and EGFP were bleached using 100% laser power, 200 iterations in a region covering around 20% of a single GUVs, and then the fluorescent signals were acquired every 2 sec. Images were analysed using ImageJ/FIJI. The resulting fluorescence was normalised to the pre-bleached images using nonbleached area as a control to correct for the recovery fluorescence and background.

### Image quantification

For **Fig. 2E** and **Fig. S3C**, GUV images were analysed using Fiji-based macro GUV-AP v2.0 (49, 50). Briefly, the GUV membranes were detected and defined by “membrane” channel, and regions of interest (ROIs) were exported as ring-like structures. The ring structure was used to quantify the GABARAP fluorescence on the membrane, while the outer circle of the ring was used to determine the total GABARAP fluorescence inside the GUVs.

### Data analysis

Statistical analysis was carried out using GraphPad Prism 10 (GraphPad Software). Differences were statistically analysed by one-way ANOVA and Tukey multiple-comparison test.

## Acknowledgments

We thank Petra Schwille (Director, Department of Cellular and Molecular Biophysics, Max Planck Institute for Biochemistry, Martinsried, Germany) for helpful technical advice and support for cDICE set-ups. We thank Terje Johansen (University of Tromso) for p62 plasmid. We thank Anne Simonsen (University of Oslo) and Alf Håkon Lystad (University of Oslo) for helpful discussion on the E3 complex purification and *in vitro* ATG8 lipidation reaction. We thank Jorge Azevedo (Universidade do Porto) for PET28-His_6_-mE1 plasmid, Katrin Rittinger for pGEX-6P1-GST-UBE2V1 and pGEX-6P1-GST-UBE2N plasmids and helpful suggestions on *in vitro* ubiquitination assay. We thank Emily Millard and Harold B.J. Jefferies (Tooze Lab, The Francis Crick Institute), Chloe Roustan in the Structural Biology facility (The Francis Crick Institute) for the technical assistance. W.Z., C.D., A.S. and S.A.T, were supported by The Francis Crick Institute which receives its core funding from Cancer Research UK (FC001187, FC001999), the Medical Research Council (FC001187, FC001999). This research was funded in whole, or in part, by the Wellcome Trust (FC001187, FC001999). W.Z. and S.A.T. received funding from the European Research Council under the European Union’s Seventh Framework Programme (FP7/2007-2013)/ERC grant agreement n° [788708]. R.D. was supported by The Francis Crick Institute, which receives its core funding from Cancer Research UK (CC1069), the UK Medical Research Council (CC1069) and the Wellcome Trust (CC1069). For the purpose of Open Access, the author has applied a CC BY public copyright licence to any Author Accepted Manuscript version arising from this submission.

## Notes

### Competing Interest Statement

The authors have declared no competing interest.

