## Supplementary figures and Table S1 for "Mechanistic studies of autophagic cargo recruitment and membrane expansion through in vitro reconstitution"

*Sharon A. Tooze

**This PDF file includes:**

Figures S1 to S5
Tables S1

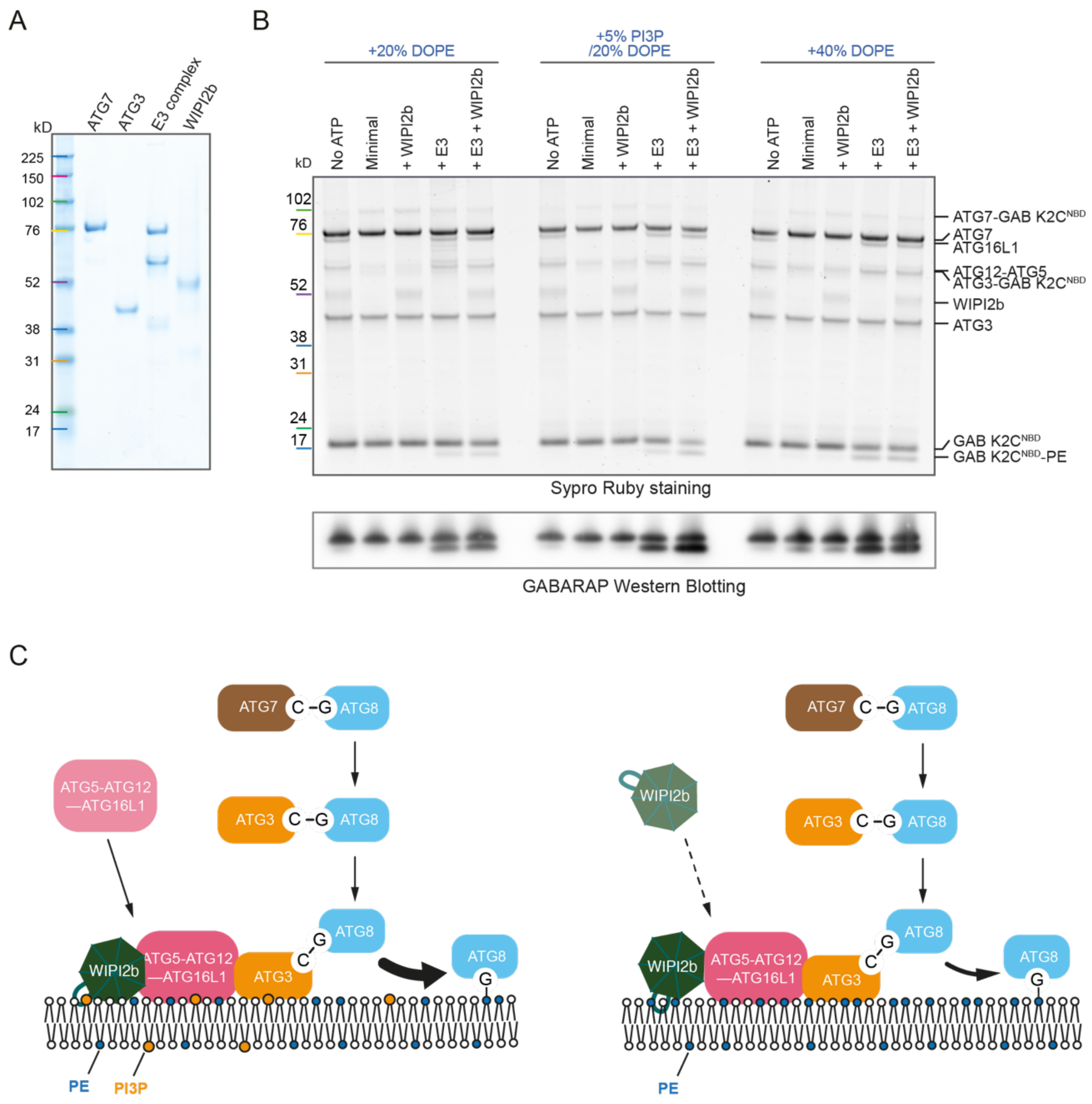

**Fig. S1 Reconstitution of WIPI2b-directed ATG8-PE conjugation machinery in vitro. (A)** SDS-PAGE of purified ATG proteins for ATG8 lipidation reaction. **(B)** SDS-PAGE and western blots of the sample after the real-time lipidation assay, shown in **Fig. 1C**. **(C)** Schematic diagram showing the dual role of WIPI2b. In the presence of PI3P, WIPI2b binds to PI3P, recruits the E3 complex to the membrane, and promote ATG8 lipidation reaction. In the absence of PI3P, WIPI2b enhances the ATG8 lipidation, compared to the E1-E2-E3 cascade alone, suggesting that WIPI2b can allosterically activate the E3 complex.

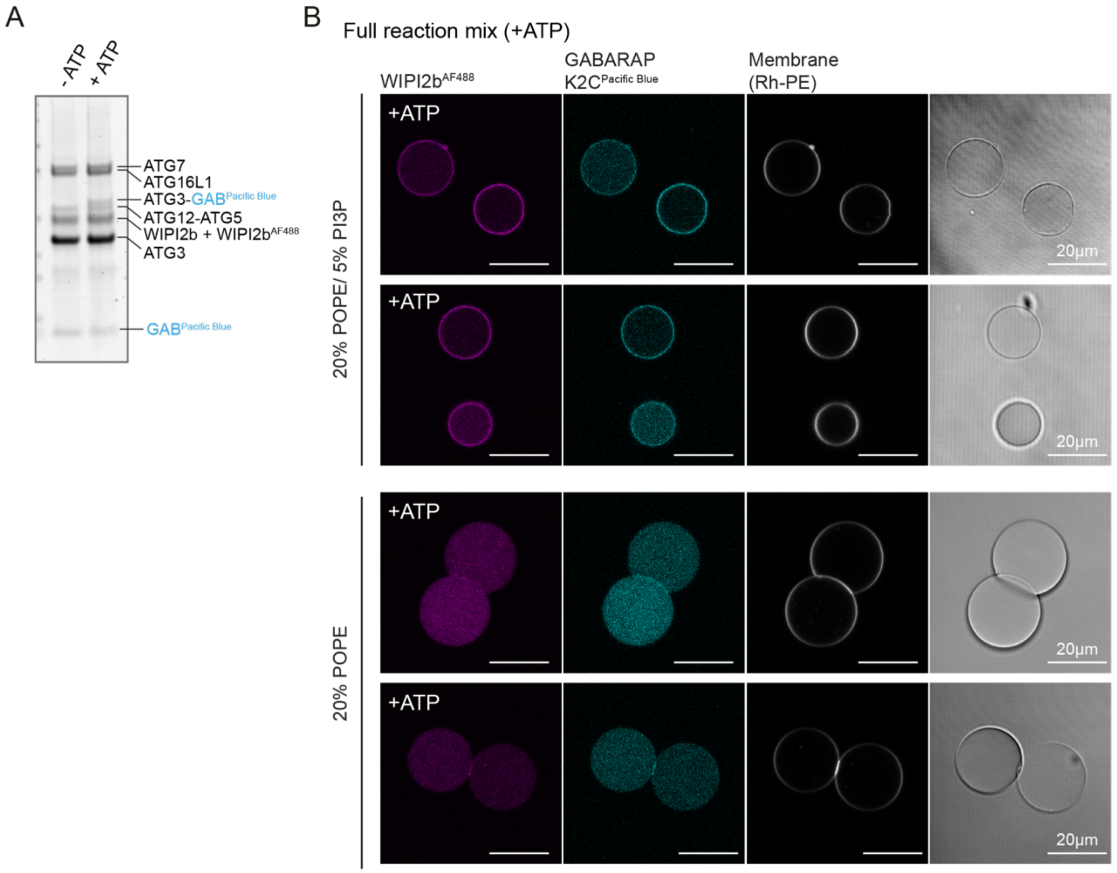

**Fig. S2. Reconstitution of WIPI2b-directed ATG8-PE conjugation reaction within GUVs. (A)** Full lipidation system with WIPI2b, with or without ATP/$MgCl_2$. The inner solution contained 0.5μM ATG7, 1.5μM ATG3, 0.5μM the E3 complex, 1.5μM WIPI2b (unlabelled to labelled at a molar ratio of 2:1), 0.5μM GABARAP K2C$^{Pacific\ Blue}$, prepared with or without 1mM ATP/$MgCl_2$. **(B)** Representative images of GUVs encapsulating the reaction mixture with ATP/$MgCl_2$. GUVs were composed of either 20% POPE/5% PI3P/74.5% POPC/0.5% Rh-PE or 20% POPE/79.5% POPC/0.5% Rh-PE.

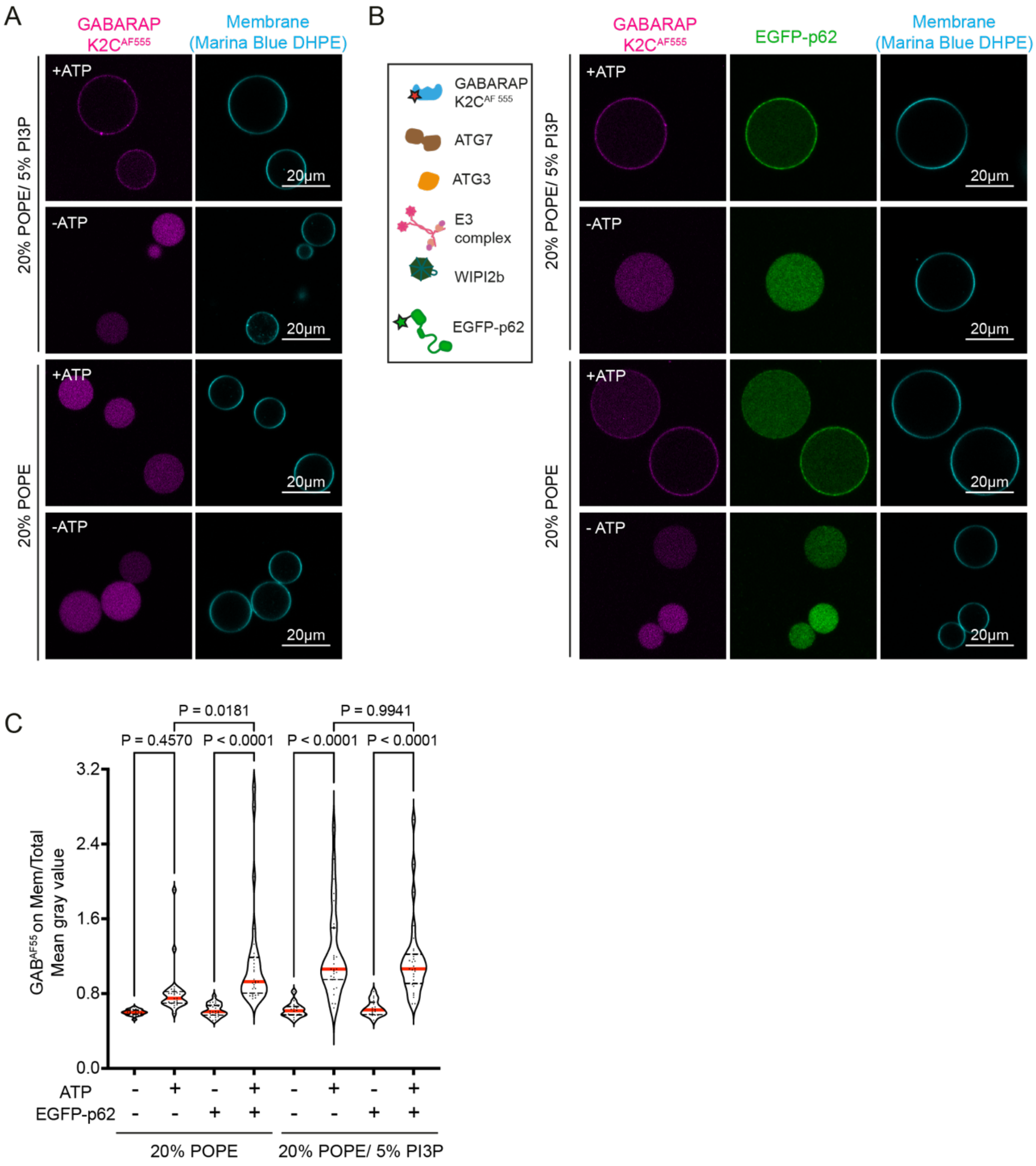

**Fig. S3 Reconstitution of cargo-directed ATG8 lipidation reaction within GUVs. (A)** Reconstitution of ATG8 lipidation inside GUVs, as described in **Fig. 2.** The inner solution contained 0.5μM ATG7, 1.5μM ATG3, 0.5μM the E3 complex, 1μM WIPI2b, 0.5μM GABARAP K2C$^{AF555}$, prepared with or without 1mM ATP/$MgCl_2$. **(B)** Reconstitution of p62-directed ATG8 lipidation reaction inside GUVs. The reaction mixture was prepared, as described in **(A)**, with addition of 1μM EGFP-p62. GUVs were composed of 20% POPE/79.5% POPC/0.5% Marina Blue-DHPE or 20% POPE/5% PI3P/74.5% POPC/0.5% Marina Blue-DHPE. **(C)** Quantification of GABARAP fluorescence on the membrane. The fluorescent intensity (mean gray value) of GABARAP on the membrane compared to that of total GABARAP encapsulated inside the GUVs (20% POPE: -ATP-p62 n=19, +ATP-p62 n=25, -ATP+p62 n=25, +ATP+p62 n=28; 20% POPE/ 5% PI3P: -ATP-p62 n=16, +ATP -p62 n=29, -ATP +p62 n=17, +ATP +p62 n=31). The thick red lines and doted black lines in the violin plot represents the medians and quartiles, respectively.

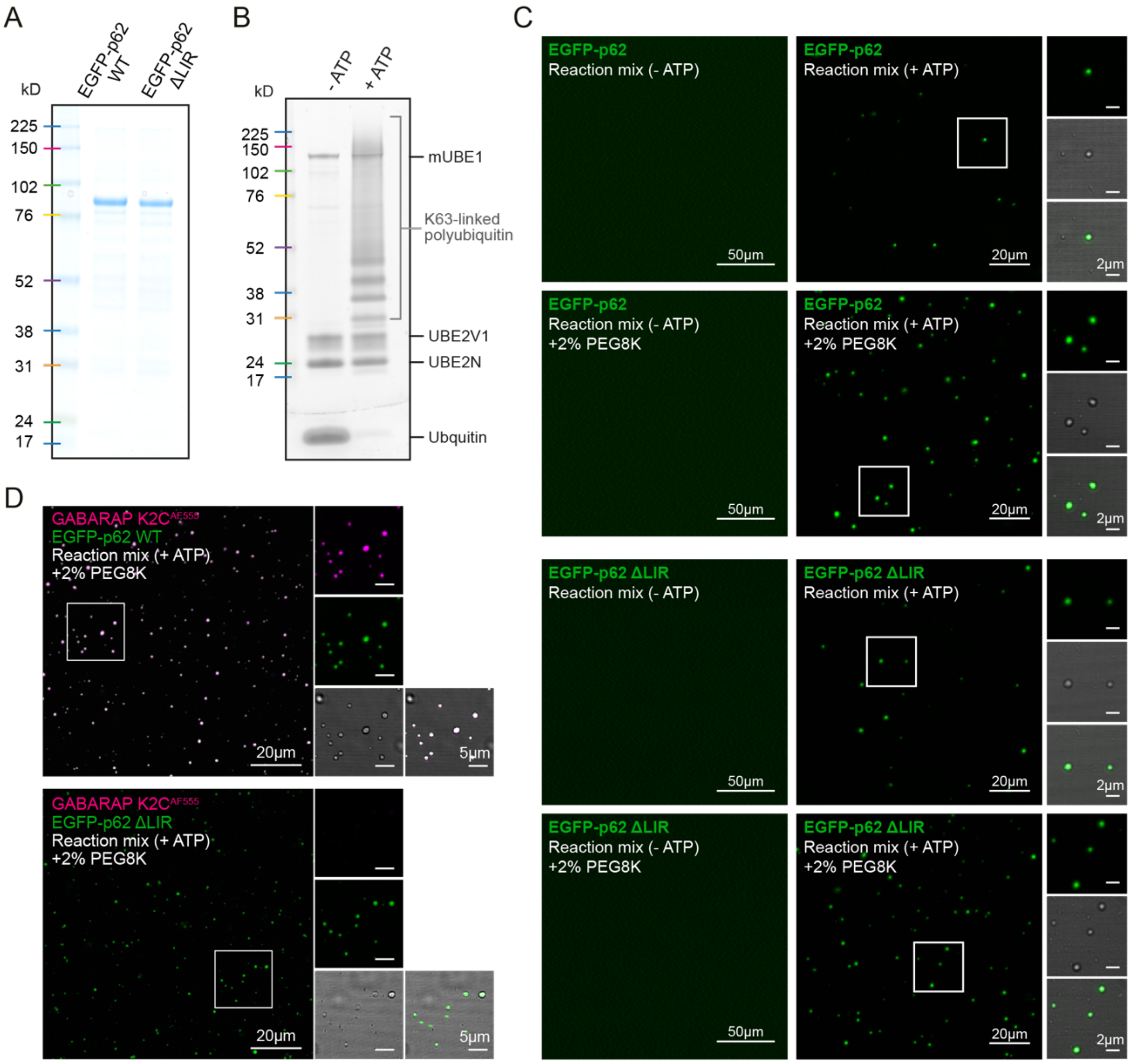

**Fig.S4 K63-linked polyubiquitin chains induce p62 phase separation. (A)** SDS-PAGE analysis of purified EGFP-p62 WT and ΔLIR. **(B)** In vitro synthesis of K63-linked polyubiquitin chains. 1μM $His_6$-mUBE1, 8μM UBE2N, 8μM UBE2V1, 100μM ubiquitin was incubated with or without 5mM ATP/$MgCl_2$ at 37℃ for 2hrs. 10μL of each reaction mix was analysed by SDS-PAGE. **(C)** In vitro reconstitution of p62 liquid-liquid phase separation. 5μM EGFP-p62 WT or ΔLIR was mixed with 20μL of the reaction mix, as described in **(B)**, with or without 2% PEG8K in a reaction volume of 100μL. **(D)** ATG8 sequestration by p62 droplets in a LIR-dependent manner. 2.5μM EGFP-p62 WT or ΔLIR and 0.5μM GABARAP K2C$^{AF555}$ was mixed with 30μL of the reaction mix (+ATP), as described in **(B)**, with 2% PEG8K in a reaction volume of 100μL. Magnified views of the outlined regions in **(C)** and **(D)** are shown in the left panels.

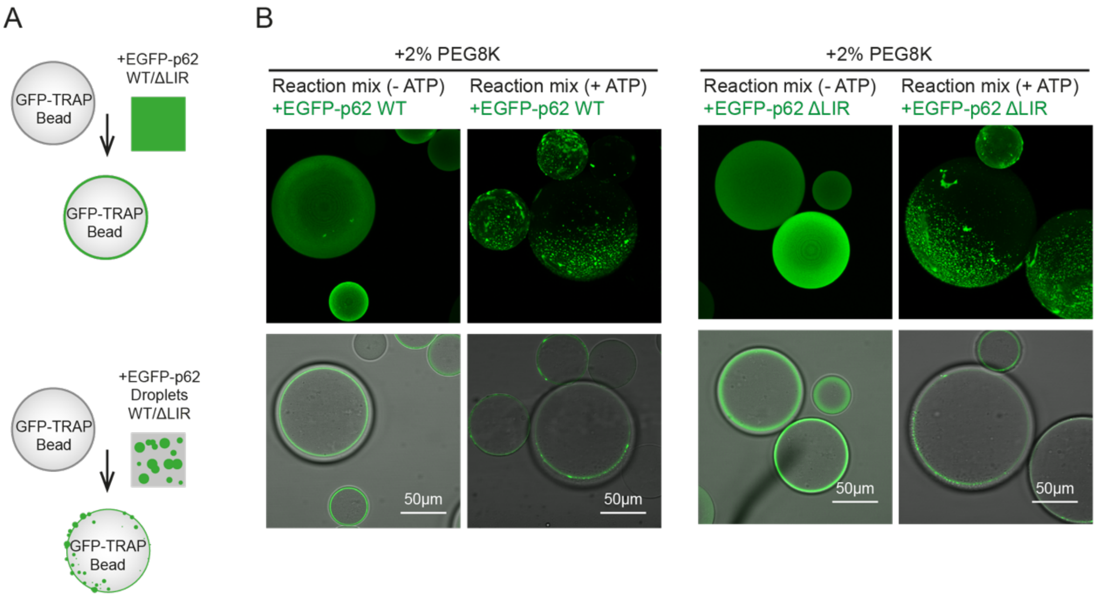

**Fig. S5 GFP-TRAP beads to generate p62 cargo template. (A)** Schematic demonstration of preparation of cargo coated beads. **(B)** Soluble or droplet-like EGFP-p62 WT or ΔLIR were coated on GFP-TRAP beads.

**Table S1.** Plasmids used in this study

| | Name | Source |
|---|---|---|
| **Bacterial expression** | pAL-GST-GABARAP$_{(1\text{-}116aa)}$ K2C | Zhang and Nishimura et al., 2023 |
| | pAL-GST-GABARAP$_{(1\text{-}116aa)}$-His$_6$ K2C | Zhang and Nishimura et al., 2023 |
| | pAL-GST-mCherry-LC3B$_{(1\text{-}120aa)}$-His$_6$ | This study |
| | pAL-GST-mTagBFP2-LC3B$_{(1\text{-}120aa)}$ | This study |
| | pGEX-6P1-GST-ATG3 | Landajuela et al., 2016 |
| | pAL-GST-EGFP-p62 | This study |
| | pAL-GST-EGFP-p62 ΔLIR | This study |
| | PET28-His$_6$-mUBE1 | Carvalho et al., 2012 |
| | pGEX-6P1-GST-UBE2V1(Uev1a) | Katrin Rittinger |
| | pGEX-6P1-GST-UBE2N(Ubc13) | Katrin Rittinger |
| **Insect cell expression** | pBacPAK-His$_3$-GST-ATG7 | Zhang and Nishimura et al., 2023 |
| | pFBDM- ATG7-ATG10-ATG12-StrepII$^{2x}$-ATG5-ATG16L1 | Zhang and Nishimura et al., 2023 |
| | pBacPAK-His$_3$-GST-WIPI2b | This study |
